# μCT-based, Three-Dimensional Cortical Bone Mineral Density Distribution reveals Unique Phenotypes related to Bone Material Properties

**DOI:** 10.1101/2025.03.18.640601

**Authors:** Serra Ucer Ozgurel, Arish Z. Maredia, Joseph Sheeran, Laurent Marichal, James C Fleet

## Abstract

Bone strength is an important factor for determining fracture risk that is dependent on bone mass and bone architecture as well as bone material properties. Bone material properties are defined, in part, by the heterogeneity and degree of bone mineralization, parameters that can be assessed with a bone mineral density distribution (BMDD) from 2D-histological sections of bone using backscatter electron microscopy (BSEM). Here we perform a comprehensive examination of a 3D-μCT-based BMDD to assess its utility in bone research. Analysis of cortical bone μCT scans from preclinical studies using anabolic treatments, pro-resorptive conditions, and genetically heterogeneous mouse lines extend and confirm published findings from BSEM-based BMDD. Principal Component Analysis identified features of the BMDD (e.g. skewness, variance, mean degree of mineralized bone or MDMB) that are distinct from the traditional bone phenotyping measures of bone mineral density, bone mineral content, and cortical bone thickness. In addition, BMDD parameters (e.g. MDMB) correlated to indices of bone material properties from Reference Point Indentation (RPI, US 1^st^, stiffness) and 4-point bending (toughness). These BMDD parameters also increased the predictive value of a multiple linear regression model for US 1^st^ from RPI (from r^2^=0.26 for traditional bone phenotypes to r^2^=0.41 for the full model). Thus, μCT-based BMDD reveals unique phenotypes related to bone material properties that complement existing bone phenotyping tools thereby increasing our ability to draw biological inferences about the nature of bone and the processes controlling bone mineralization.

## Introduction

In addition to bone mass and cortical/trabecular bone microarchitecture, the material Properties of bone (MP) are an important factor for determining bone structural integrity (strength) and fracture risk.^1,2^ However, classical bone assessment tools aren’t sufficient predictors of bone strength or osteoporosis risk. The gold standard for assessing bone health is bone mineral density (BMD), an index of bone mass measured by Dual Energy X-ray Absorptiometry (DEXA). However, in a meta-regression analysis of 38 clinical trials, the change in BMD due to treatment predicts just 48% of hip fracture risk.^3^ A second independent predictor of fracture risk is cortical and trabecular bone microarchitecture, which is assessed by high resolution-peripheral quantitative computational tomography (HR-pQCT) or micro-computational tomography (μCT).^4^ In contrast, MP are determined by various aspects of bone tissue including collagen content and crosslinking in bone, microdamage to bone, and mineralization.^5^ Mineralization can be broken into two major parameters: the degree and the heterogeneity of bone mineralization. However, heterogeneity of mineralization is not measured by typical bone imaging tools; DEXA reduces the three-dimensions of bone into a two-dimensional image while HR-pQCT and μCT reduce the bone mineral levels to binary bone/not-bone feature that masks the heterogeneity.

Bone mineralization can be quantified by several methods. In laboratory settings, the most common procedure uses dynamic histomorphometry. While this approach is useful for determining the rate of new bone mineralization, it does not determine the degree or heterogeneity of the bone. Another procedure quantifies a bone mineral density distribution (BMDD) reflecting the degree and heterogeneity of bone using backscatter electron microscopy (BSEM)^6^. This approach can be applied to clinical populations but is limited by the requirement for a bone biopsy and the need for specialized equipment. While some studies have used μCT to generate a BMDD^7–10^, a comprehensive examination of μCT-based BMDD has not yet been conducted.

To this end, our research had two major objectives. First, we assessed whether μCT-based BMDD can be reliably used for assessing degree and heterogeneity of bone mineralization. For this objective, we analyzed cortical bone samples from preclinical studies using a variety of experimental conditions known to alter bone mass, e.g. anabolic treatments, pro-resorptive conditions, genetically distinct mouse lines, and environmental challenges like low calcium intake. Second, we compared BMDD parameters to traditional bone measures (e.g. BMD, bone mineral content (BMC), cortical μCT parameters) to identify features of the BMDD that correlate uniquely to indices of MP. Our research reveals that μCT-based BMDD is easily applicable and provides unique phenotypes related to MP that complement existing bone phenotyping tools.

## Results

### 1.1. Impact of Pro-resorptive Interventions on BMDD

The pro-resorptive actions of osteoclasts are proposed to modify the BMDD by driving the removal of high density tissue in favor of newly formed lower mineral densities.^11^ As our initial test of the μCT-based BMDD method, we chose an extreme example of under mineralization - osteomalacia due to abnormal calcium metabolism in Vitamin D knockout (VDR KO) mice.^12^ VDR KO mice had a dramatically different BMDD phenotype compared to littermate wild type (WT) mice (Fig. 1A&B). They had less total bone (-34.1% total number of voxels, -62.9% height, Supplemental Fig. 2, Supplemental Table 1) and most of the bone was represented as lower grayscale values (i.e. low mineralization, Fig. 1B). The shift to lower density in VDR KO was reflected in a lower MDMB (Fig. 1C), center, lower 5%, upper 5%, and Max values compared to WT mice (Supplemental Table 1). VDR KO also changed the shape of the BMDD as reflected in a less negative skewness and less leptokurtic kurtosis (i.e. indicating a broader BMDD). Consistent with the lower kurtosis value, FWHM was higher in VDR KO, indicating there was a more heterogenous BMDD compared to WT.

**Figure 1.**
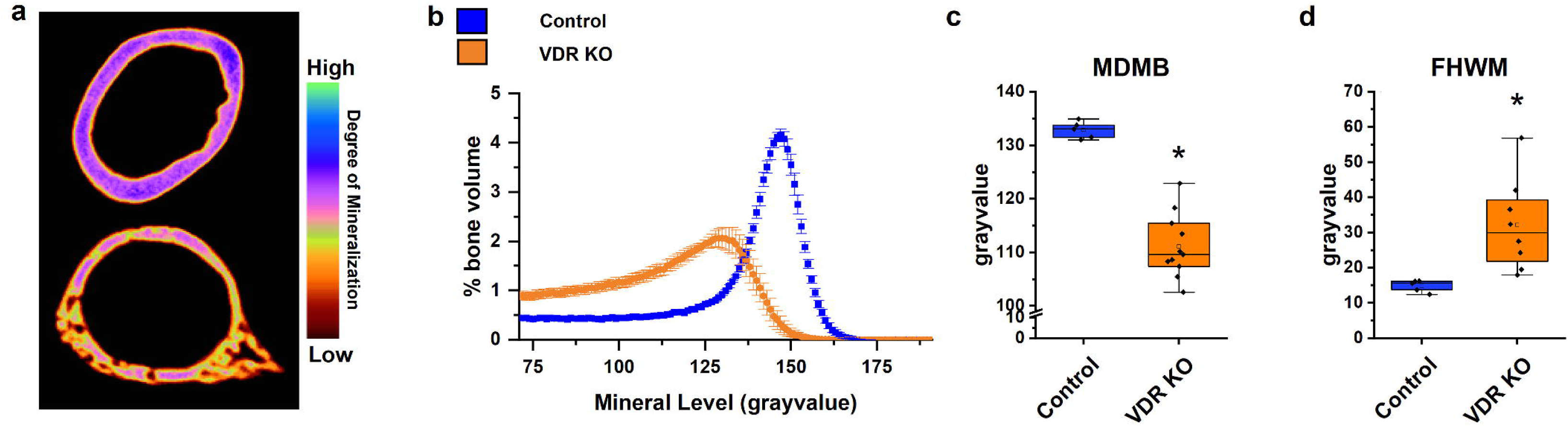
The Impact of VDR KO on Femoral Midshaft BMDD: **A.** Representative false color image of μCT data demonstrating the BMDD of femoral cortical midshaft in 12-wk old vitamin D receptor knockout (VDR KO n=12) and littermate control (n=5) mice fed 0.5% Ca AIN93G diets. Colors depict the degree of mineralization. Images are generated utilizing DataViewer. **B.** Normalized BMDD curve. BMDD is plotted as mean±SEM., **C.** MDMB and **D.** FWHM from BMDD (n=8 VDR KO). Dot plots are superimposed over box plots with the central box spanning the 25th and 75th percentiles, the central line representing the median, and the whiskers representing the 5^th^ and 95^th^ percentiles. *p<0.05 Analyzed by t-test.

To further evaluate the pro-resorptive actions of osteoclasts on bone, we chose two additional models: disease-based bone loss induced by chronic kidney disease (CKD) and the cumulative bone loss due to aging. Consistent with a previous study that reported both CKD and aging promoted cortical thinning and reduced cortical bone area,^13^ we observed that in both cases, there was less total bone (e.g. peak height: CKD=-40.5%, aging=-24.8%, Fig. 2A&B, Supplemental Table 1) while the MDMB and lower 5% cutoff of the BMDD were significantly reduced. The lower mineralization in CKD and aging also changed the shape of the BMDD; the distribution shape changed from leptokurtic to normal and became less left skewed. Finally, variance of the BMDD increased indicating greater heterogeneity in both groups.

**Figure 2.**
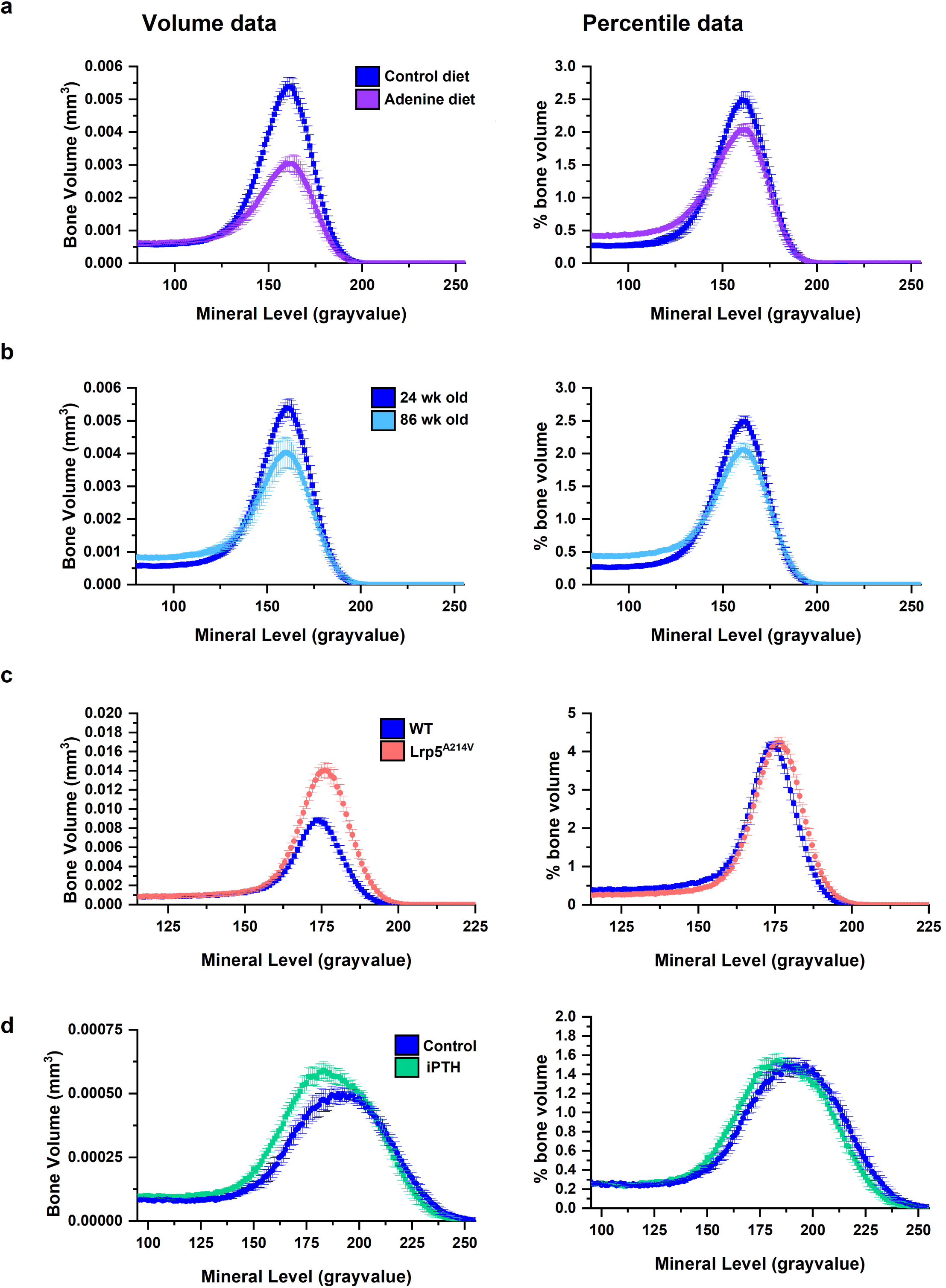
Impact of Pro-resorptive and Anabolic Conditions on BMDD: Volume (left) and normalized (right) BMDD of femoral cortical bone from **A.** 24-week-old male mice with adenine diet inducing CKD (n=7) or control diet with normal kidney function (n=7); **B.** 24-week (adult, n=7) or 86-week-old (old, n=7) male mice; **C.** 12 wk-old Lrp5^A214V^ (n=9) and their littermate control (n=9) male mice. **D.** 22-week-old male mice treated with vehicle (n=7) or PTH 1-34 (100 ng/g/d) (n=7) for 28 days. BMDD data are plotted as mean±SEM.

### 1.2. Impact of Anabolic Interventions on BMDD

The anabolic actions of osteoblasts are proposed to modify the BMDD by driving the distribution to higher mineral densities.^11^ We evaluated the anabolic actions mediated by osteoblasts and osteocytes by examining the impact of the Low-density lipoprotein receptor-related protein 5 with gain of function gene mutation (Lrp5^A214V^)^14^ and of intermittent PTH (iPTH) administration^15^ on the BMDD.

Lrp5^A214V^ mice have higher cortical bone mass and % ash weight due to continuous, high-level osteoblast activity.^16–18^ Consistent with the higher bone mass, we saw a 55.2% increase in the number of voxels and a 57% increase in peak height (Fig. 2C). In addition, the upper and lower 5% cutoffs, the MDMB, and the Maximum gray value of the BMDD were shifted upwards in Lrp5^A214V^ mice. This is consistent with the more negative skewness to the BMDD we observed in Lrp5^A214V^ mice and indicates a higher degree of mineralization. The Lrp5^A214V^ BMDD also had a more leptokurtic shape and an increased FWHM that indicated that although the bone was more mineralized it was also more heterogeneous.

Mice treated daily with iPTH for 28 days had more bone (+14.7% voxel #, +21.2% height, Supplemental Table 1). While the Maximum gray value was similar in both iPTH and control groups, the new bone added in the iPTH group was to the lower greyscale part of the BMDD and both the upper 5% cutoff and MDMB were lower, reflecting lower mineralization levels (Fig. 2D). Kurtosis, FWHM, and skewness were similar in the two groups demonstrating that the overall shape of distribution was not changed other than shifting the distribution of mineralization to a lower level with iPTH. Overall, this suggests that the newly added bone resulting from 28 d of iPTH treatment did not have enough time to go through secondary mineralization.

### 1.3. Genetic Diversity Impacts the BMDD

Another biologically important factor that controls bone is natural genetic variation. We have previously shown the importance of genetics as a modulator of traditional bone phenotypes from DEXA and μCT using a panel of 11 inbred mouse strains chosen to capture >90% of the genetic variation in the mouse genome.^19^ Like our previous report, there were significant line (i.e. genetic) effects influencing all 11 BMDD parameters (Fig. 3A, Supplemental Table 3). Like with BMD, there were lines with high (AKR/J, C3H/HeJ, 129S1/SV) and low (PWK, CAST, WSB) levels of the BMDD parameters that reflect bone mass (i.e. number of voxels, peak height)Fig.. There were also different patterns of skewness (C3H/HeJ, 129S1/SV, and AKR/J being more left skewed and C57BL/6J, PWK/PhJ, and SWR/J being more symmetrical) and kurtosis (C3H/HeJ, AKR/J, CBA/J being more leptokurtic and C57BL/6J, PWK/PhJ, and SWR/J being more normally distributed). Two parameters reflecting heterogeneity were also different across the lines, i.e. FWHM (DBA/2J, PWK/PhJ, and C57BL/6J being wider while SWR/J, AKR/J, and 129S1/SV being narrower) and variance (DBA/2J, WSB/EiJ, and CBA/J being more variable while SWR/J, C57BL/6J, and C3H/HeJ being less variable). As a preliminary test to validate that we are not simply assessing bone mass with the BMDD parameters, we compared the strain distribution of femoral BMD with MDMB across the 11 lines. Fig. 3B is organized from the line with the highest (CBA/J) to lowest (C57BL/6) MDMB level and shows that the strain distribution for BMD did not follow the same pattern as for MDMB (e.g. values for DBA/2J, CAST/EiJ, WSB/EiJ, and A/J are particularly divergent).

**Figure 3.**
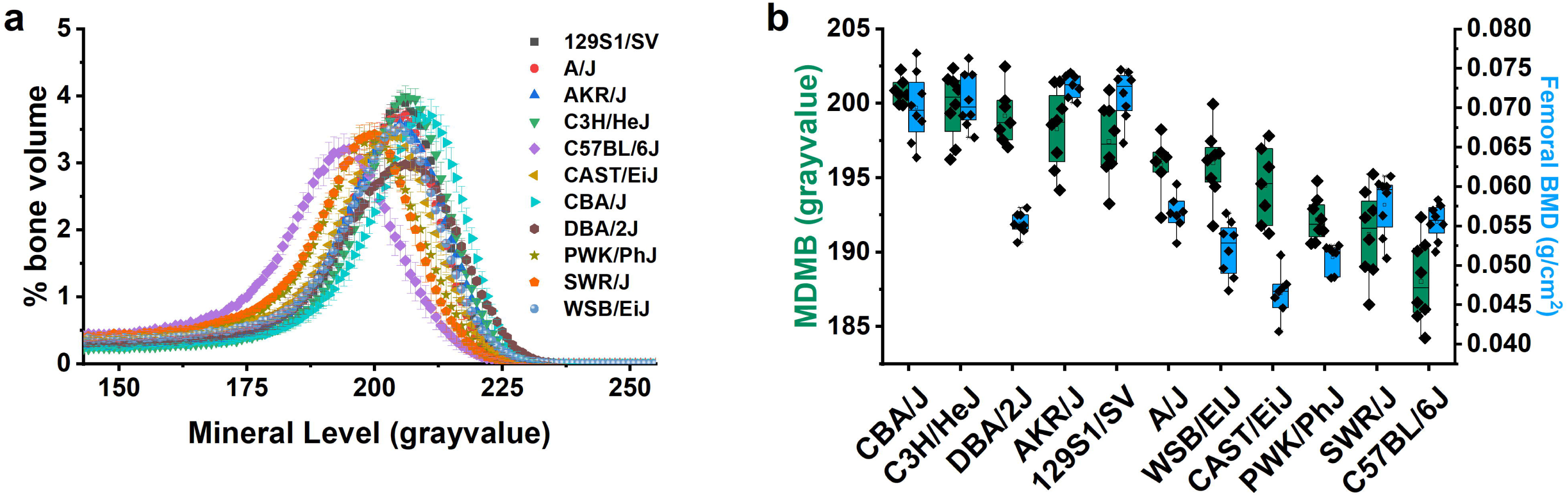
The Impact of Natural Genetic Variation on the Femoral BMDD. **A.** Normalized BMDD of cortical femur midshaft of 12-week-old male mice from 11 inbred strains fed 0.5% Ca diets (n=6-8/group). BMDD data are plotted as mean±SEM. B. The strain distribution of MDMB and femoral BMD of the mice from A (n=6-8). Dot plots are superimposed over box plots with the central box spanning the 25th and 75th percentiles, the central line representing the median, and the whiskers representing the 5^th^ and 95^th^ percentiles.

### 1.4. Low Calcium Diet Modified BMDD

Classically, dietary calcium restriction causes physiologic adaptation leading to elevated serum PTH and 1,25(OH)_2_D levels that increase the efficiency of intestinal calcium absorption and renal calcium reabsorption. However, if this adaptation is not sufficient to compensate for the loss of dietary calcium, bone resorption will increase and bone mass will decline.^20^ Across the 11 inbred lines, low calcium intake reduced BMC, the number of voxels, peak height, the lower 5% cut off, and made the distribution less leptokurotic. However, the response was highly variable across the lines; in six lines the BMDD parameters were unchanged, while in five lines the effect of diet was seen on multiple BMDD parameters (Supplemental Table 3). One example of this diversity is the impact that low calcium has on skewness; while this parameter is unchanged in C3H mice, in CAST mice the distrbution became more symetrical and more normally distributed (Fig. 4C,D). As we previously reported^19^, BMD is altered by low calcium intake in some strains (AKR, C3H, CBA), but not the others, indicating the existance of gene-by-diet interaction (Fig. 4A). The strain distribution patterns showed that low calcium diets had a negative effect on both BMD and skewness, but theeffect on skewness was observed in different lines than for BMD (higher in AKR, 129S1, DBA, CAST, Fig. 4B).

**Figure 4.**
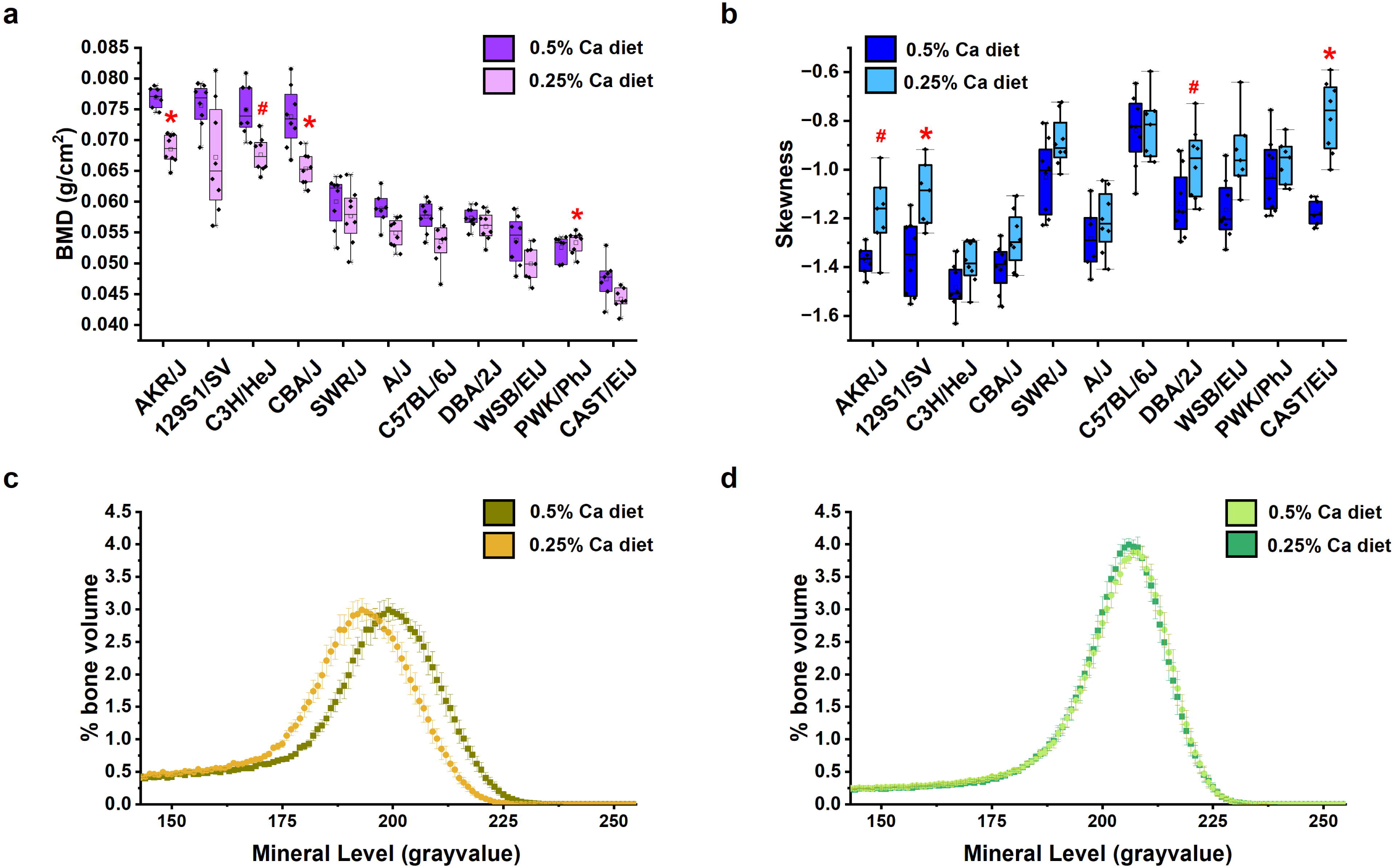
Gene-by-Diet Effects on BMDD. The strain distribution of **A.** femoral BMD and **B.** femoral cortical midshaft BMDD skewness from of 11 inbred strains of male mice fed either 0.25% or 0.5% Ca diet from 4-12 weeks of age or (n=6-8/diet). Dot plots are superimposed over box plots with the central box spanning the 25th and 75th percentiles, the central line representing the median, and the whiskers representing the 5^th^ and 95^th^ percentiles. *p<0.05, #p<0.10 vs control diet group within an inbred line. Normalized BMDD data from **C.** CAST/EiJ mice and **D.** C3H/HeJ mice fed 0.25% or 0.5% Ca diets (n=7-8/diet). BMDD data are plotted as mean±SEM.

### 1.5. Relationships between BMDD Parameters and Traditional Bone Phenotypes

We conducted several analyses to further test how BMDD phenotypes relate to traditional bone phenotypes. As expected, simple linear regressions show that the number of voxels and peak height were strongly correlated to BMD, BMC, and Cortical thickness (Ct.Th) (e.g. r^2^=0.859 and 0.774 to BMC, respectively, Supplemental Table 4). A strong relationship was also found between Ct.Th and the lower 5% cutoff (r^2^=0.869) while more moderate relationships were found between Ct.Th and kurtosis (r^2^=0.736), and skewness (r^2^=0.517). Weaker relationships were found for the other BMDD parameters with traditional bone phenotypes (e.g. r^2^=0.375 for skewness and BMD: 0.255 for MDMB and BMD; -0.211 for variance and BMC; 0.057 for FWHM and BMD). Thus, while a portion of BMDD parameters is predicting similar biology as traditional bone measures, they reflect unique aspects as well.

To further clarify the novelty of the BMDD parameters, we used Principal Component Analysis (PCA) to identify the linear combinations of traditional bone parameters and BMDD phenotypes that capture unique aspects of the variance controlling bone biology. The Scree plot revealed three principal components (PC) accounting for 92.2% of the variance in the dataset. PC1 with 59.1% of variance was driven by the traditional bone phenotypes (BMD, BMC, Ct.Th) and the BMDD parameters correlated to them (i.e. # voxels, peak height, lower 5% cutoff) as well as MDMB, kurtosis, and skewness. PC2 (23.8%) was driven by the variance, upper 5% cutoff, Max grayvalue, center, and MDMB while PC3 (9.24%) was driven by FWHM, skewness, and Max grayvalue (Fig. 6 and Supplemental Fig. 5). These three distinct and independent predictors separate clearly when visualized in a 3D loading plot (Fig. 6).

**Figure 5.**
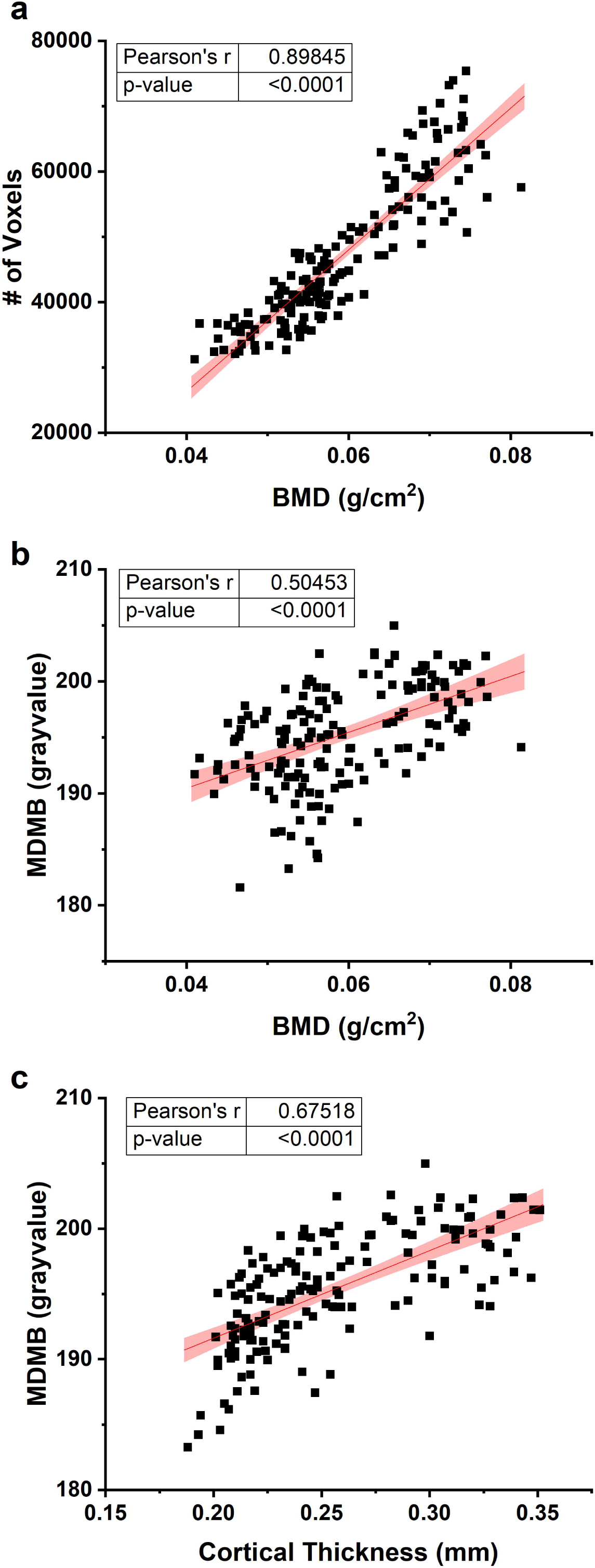
Correlation of BMDD Parameters with Traditional Bone Phenotypes: Pearson’s correlations were performed between femoral BMD (obtained from DEXA) and **A.** femoral cortical midshaft number (#) of voxels. (n=167) or **B.** MDMB (n=165); or **C.** Cortical Thickness and MDMB from 11 genetically diverse inbred mouse strains fed 0.25% and 0.5% Ca from weaning to 12 week of age (n=159). The solid red line is the linear regression line and the pink are is 95% confidence interval of the data. Pearson’s r and corresponded p-value for each graph are reported in tables within the graphs.

**Figure 6.**
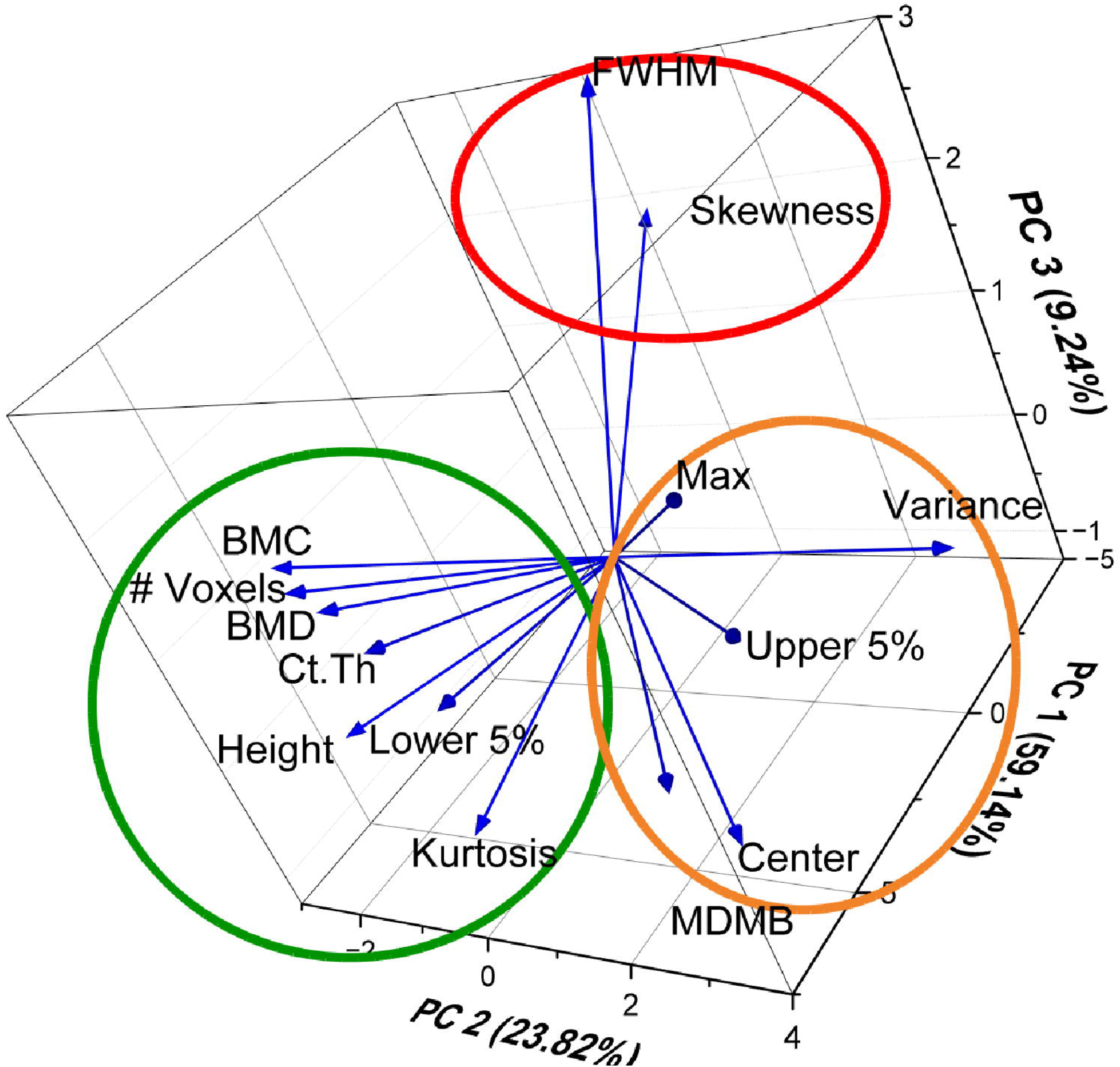
PCA Analysis Reveals Different Categories of BMDD Parameters: A 3D PCA loading plot shows some BMDD parameters correspond to traditional bone phenotypes (green circle) while others are unique descriptors of bone (red, orange circles).

### 1.6. BMDD Correlates with MP

DEXA, μCT, and MP each provide data that contribute to the structural integrity of bone but mechanical testing with 3- or 4-point bending or with reference point indentation (RPI) are more direct measures of bone mechanical properties.^21,22^ Initially, we used Pearson’s correlations to determine the relationship between femoral BMDD parameters or traditional bone phenotyping tools and RPI in a population with a large degree of heterogeneity due to genetic (11 inbred mouse lines) and dietary (low vs adequate calcium) factors (Fig. 7A, 7B; Supplemental Table 4). The strongest Pearson’s correlation coefficients were seen between US 1^st^ (a measure of bone material stiffness)^23^ and this was true for traditional bone assessment tools (e.g. BMD, r=0.49; Fig. 7A) and BMDD parameters (e.g. MDMB, r=0.548; Fig. 7B). We also conducted Pearson’s correlation analysis between data from 4-point bending and BMDD parameters from the CKD and aging study (Supplemental Table 5). Consistent with the other analysis, we found strong correlations between many BMDD parameters and stiffness (e.g. MDMB, r=0.629). In addition, we saw that many BMDD parameters were also correlated with toughness, (e.g. MDMB, r=0.511; Fig. 7C).

**Figure 7.**
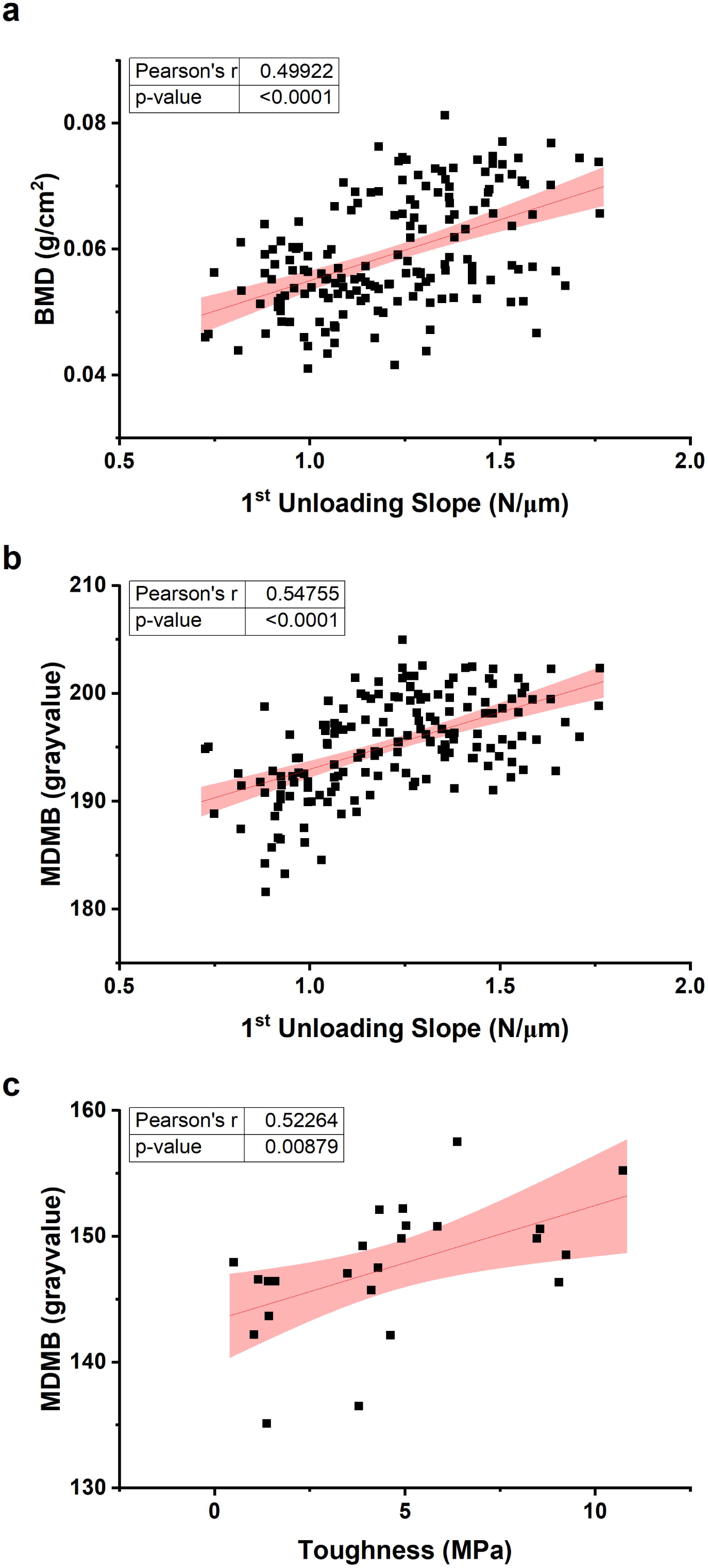
BMDD Parameters Correlate with Material Properties of Bone: Pearson’s correlations were performed between **A.** femoral BMD and 1^st^ unloading slope, a measure of bone stiffness. (n=168) or **B.** MDMB and 1^st^ unloading slope (n=166) from 11 genetically diverse inbred mouse strains fed 0.25% and 0.5% Ca from weaning to 12 week of age. **C.** MDMB and bone toughness from adenine or control fed young and old mice. (n=24). The solid red line is the linear regression line and the pink are is 95% confidence interval of the data. Pearson’s r and corresponded p-value for each graph are reported in tables within the graphs.

Because our PCA analysis indicates that traditional and BMDD parameters have independent features, we further examined how these multiple independent variables influence bone stiffness. We used a backward selection multiple linear regression model using three different phenotype combinations: the three traditional bone phenotyping parameters, the 11 BMDD parameters, and a combination of the traditional and BMDD parameters (14 variables) (Table 1). The traditional parameters model and the BMDD parameters improved the correlation with US 1^st^ compared to the best individual Pearson’s correlation in each group (i.e. BMD, r=0.499; MDMB, r=0.548). In addition, the BMDD model had a stronger correlation with US 1^st^ that the traditional bone parameters model (r=0.597 vs 0.52). However, the strongest correlation was seen for the combined model (r=0.689 accounting for 41.3% of the US 1^st^ phenotype variance) due to contributions from both traditional bone phenotypes (BMC, Ct.Th) and BMDD parameters (# voxels, FWHM). Overall, this analysis demonstrates that including BMDD parameters in the models improved the ability to predict bone stiffness, an important indicator of the MP.

**Table 1.** Multiple Linear Regression to Predict the 1^st^ Unloading Slope from Traditional and BMDD Parameters.

| Parameters | # variables | R-value | Adj R<sup>2</sup> * | n | df | Significant Parameters |
| --- | --- | --- | --- | --- | --- | --- |
| BMD, BMC, CtTh | 3 | 0.520 | 0.256 | 154 | 140 | CtTh, BMD<sup>#</sup> |
| All BMDD | 11 | 0.597 | 0.305 | 149 | 137 | FWHM, MDMB<sup>#</sup> |
| All (BMD, BMC, CtTh, BMDD) | 14 | 0.689 | 0.413 | 133 | 118 | BMC, total voxels, CtTh<sup>#</sup>, FWHM<sup>#</sup> |
| *\* Adjusted for the number of independent variables included in the model* |  |  |  |  |  |  |
| <sup>#</sup> p<0.08 |  |  |  |  |  |  |

## Discussion

MP is one of three critical parameters, along with bone mass and bone geometry/architecture, that determine bone structural integrity.^24^. The BMDD measured by BSEM or other methods (e.g. μCT, microradiography) carries valuable information about MP.^25,26^ For example, a BMDD quantifies the heterogeneity of mineralization, a parameter that limits microcrack propagation^27^ and influences bone strength.^28^ In addition, the BMDD reflects the balance between osteoblast/osteocyte-mediated formation/mineralization and osteoclast mediated resorption events and so it is viewed as a fingerprint of mineralization.^11^ However, because BSEM and other methods are cumbersome, BMDD is not commonly measured in the clinic or in research studies. Here, we report a careful, comprehensive assessment of BMDD derived from μCT. To achieve this goal, we used a wide variety of preclinical phenotypes to fully capture the potential of the method for assessing the impact of physiologic, genetic, or pharmacologic changes on cortical BMDD. We have shown that μCT-based BMDD measures features of bone mass (i.e. consistent with BMD, BMC) but also reveals district measures of bone heterogeneity determined from the shape of the distribution (i.e. variance, FWHM, kurtosis, and skewness), and this information enhances prediction of bone mechanical properties beyond what can be achieved with traditional bone phenotypes obtained from DEXA (BMD) or μCT (Ct.Th). Overall, this shows that utilizing μCT-based BMDD is a simple and feasible approach to capturing unique features that contribute to the MP.

We selected various preclinical mouse models to determine if μCT-based BMDD can recapitulate and extend the previous knowledge about mineralization. Previously, Roschger *et al.*^29^ assessed BMDD of a transiliac bone biopsy from a patient with osteomalacia and found that the bone was more heterogenous (quantified by FWHM) and less mineralized (quantified as Ca_mean_, a weighted average calcium concentration similar to MDMB). Here we used μCT-based BMDD to evaluate the osteomalacia that develops in VDR KO mice due to elevated osteoclast activity resulting from severe disruption of calcium metabolism.^12^ In agreement with the patient data, the mouse model exhibited lower bone mass (height, total number of voxels) as well as a severely left skewed, platykurtic distribution and a larger FWHM reflecting a high volume of under mineralized bone and increased heterogeneity, respectively.

We next sought to evaluate BMDD in two bone catabolic conditions with increased osteoclast activity that promoted either acute (CKD) or chronic (aging) bone loss.^30,31^ Misof *et al.*^32^ previously used BSEM-based BMDD on iliac crest biopsies from 58 CKD patients who were classified into high/normal/low turnover groups as determined by histomorphometry. BMDD from both the high and low turnover groups had a lower mean degree and peak bone mineral as well as higher heterogeneity (FWHM) than the non-CKD group. While we did not measure bone turnover rate in our CKD mouse model, our results are like those of human subjects with low turnover, i.e. less total bone (lower total number of voxels and height) and more heterogeneity (less leptokurtic distribution and higher variance). Roschger *et al.*^29^ found no correlations between age and BSEM-based BMDD parameters (i.e. peak, mean, and FWHM) in a cross-sectional analysis of people between 25 and 97 years-old. However, with more BMDD parameters in our tightly controlled experiment, we showed that aging caused the BMDD to become more heterogeneous (i.e. less leptokurtic and more right-skewed distribution with higher variance) with less mineral (lower MDMB and lower 5% cutoff). Our data on aging is also in agreement with the mathematical calculations of bone remodeling from BSEM-based BMDD by Ruffino *et al.*^11^, which demonstrated that increased osteoclast activity decreases kurtosis.

To assess the utility of μCT-based BMDD for evaluating anabolic conditions on bone, we selected two animal models with high osteoblast activity that was constitutive (Lrp5^A214V^) or acute (iPTH). Research shows that Lrp5 gene mutations cause chronic elevations in osteoblast Wnt signaling that lead to higher bone mass in humans and mice.^33^ Using BSEM-based BMDD, Roetzger *et al.*^34^ demonstrated that a patient with a Lrp5-activating gene mutation had higher mineralization and less heterogeneity in skull cortical bone. Furthermore, Ross *et al.*^18^ demonstrated that Lrp5^G171V^ mice had increased femoral bone strength (assessed by 3-point bending) and an increased mineral-to-matrix ratio indicative of a more highly mineralized, higher quality bone. Our μCT-based BMDD data are consistent with these studies by showing that Lrp5^A214V^ mice have higher bone mass (height and voxel #), with higher peak, MDMB, and maximum grayvalue (indirect indicators of improved MP) and less heterogeneity (kurtosis, skewness, and variance). iPTH treatment is a FDA-approved treatment for osteoporosis^35^ that stimulates osteoblast activity, leading to production of new bone in humans and rodents.^36^ Using BSEM-based BMDD Misof *et al*.^37^ demonstrated that following 18 months of iPTH treatment, iliac crest biopsies of osteoporotic men exhibited a reduced MDMB without changes in heterogeneity. In addition, they found that after iPTH treatment there were significant correlations between histomorphometric parameters of bone turnover and the BMDD parameters Ca_Peak_ and FWHM (r^2^=0.46 and 0.45 for osteoid perimeter; r^2^=0.63 and 0.97 for BFR). Their data suggests that under-mineralized bone can be revealed by BSEM parameters, and they concluded that the lower degree of mineralization observed following iPTH treatment is transient and will reach to a mature state during secondary mineralization. In agreement with their data, we found that although there was more cortical bone in male mice treated with iPTH for 4 weeks, the newly added bone was added to the lower mineralization level of the BMDD. Thus, this data is important for understanding the process of bone mineralization since the μCT-based BMDD method can also reveal when mineralization is not yet fully consolidated.

In addition to catabolic and anabolic conditions, bone phenotype variation in mice and humans can be driven by natural genetic variation.^38^ In humans, the impact of genetics on BSEM-based BMDD only been examined in one small study by Roschger *et al.*^29^ They found no differences in mean, peak, or FWHM in the BSEM-based BMDD of iliac crest biopsies from African American (n=15) and Caucasian (n=27) premenopausal women. In contrast, we revealed that genetic diversity is a strong driver of BMDD. All 11 BMDD parameters we measured were significantly affected across the genetically diverse population of 11 inbred mouse lines we assessed (e.g. min/max differences among lines were high for bone mass measures (total # voxels +100%, peak height +122.2%) as well as heterogeneity parameters: (variance +24.48%, kurtosis +53.7%). This is likely due to the fact we captured more genetic diversity in our well-controlled study with 85 age- and sex-matched mice from 11 inbred mouse lines. Our observations from these mice open the potential to use a forward genetic mapping approach to find variation in genes controlling the degree and heterogeneity of bone mineralization.

BMDD provides unique features of MP that contribute bone structural integrity, i.e. the degree and heterogeneity of bone mineral^5^, so we next evaluated the relationship of our μCT-based BMDD to indices of bone mechanical properties. Two papers previously reported relationships between BMDD parameters from bone tissue biopsies and indices of MP. Follet *et al.*^26^ found that in calcanei from 20 elderly subjects, MDMB (similar to DMB in the paper) was correlated with elastic modulus (r^2^=0.44) and maximum strength (r^2^=0.41); they concluded that higher MDMB is an indicator of stiffer bones. Hoc *et al.*^25^ examined the relationship between BSEM-based BMDD femoral cortical bone biopsies and MP from nanoindentation of a single cow femur and found that MDMB was significantly correlated with Young’s elastic modulus (r^2^=0.75). Although these papers show the importance of the degree of mineralization as a surrogate for MP, both use the 2D BSEM-based BMDD analysis and neither examined all of the features that define a BMDD. Here we extend these findings by comparing μCT-based BMDD parameters to MP from various mechanical tests using data from two controlled rodent studies. In our large, genetically diverse population of mice, we found significant correlations between MDMB and the RPI-derived measure of material stiffness (US 1^st^). Additionally, we found significant correlations between MDMB and toughness performed in CDK and aging study using 4-point bending test. Importantly, we found with multiple linear regression modeling that BMDD parameters like MDMB and FWHM added additional predictive value for US 1^st^ beyond that provided by traditional bone phenotypes like BMD, BMC, and Ct.Th. These data both confirm earlier relationships defined with 2D methods and extend the validity of the 3D μCT-based method as a surrogate measure of MP.

The primary strength of this work is that we established that μCT-based BMDD works in translational as well as in physiological settings and we showed that it recapitulates BSEM-based BMDD results. This was accomplished using a diverse array of existing datasets that allowed us to demonstrate that these new parameters of bone mineralization are biologically relevant. In doing so, our data significantly extended previous work on μCT-based BMDD by others. In an earlier analysis, Mashiatulla *et al*.^7^ used a panel of HA standards and a variety of single bone samples from 8 different species to compare μCT-based BMDD to BSEM-based BMDD. While they found μCT- and BSEM-based BMDD correlated for a number of BMDD parameters, they did not utilize the method to assess any physiological or disease states. Later, Tu *et al,*^8^ showed that dietary magnesium deficiency disturbed BMDD parameters from mouse femur cortical bone midshafts. However, while they explained the BMDD parameters they measured, they did not present any data to validate the methodology. Subsequently, Parle *et al.*^9^ used μCT-based BMDD on cancellous bone core samples from femoral heads to demonstrate differences in mechanical features (strength, stiffness, loading energy, using a single-column tensile/compression testing machine) and BMDD parameters (MDMB, Peak center, FWHM) among samples from patients with osteoarthritis, osteoarthritis with type 2 diabetes, and osteoporosis. Yet they didn’t correlate the BMDD values to the MP indices. Finally, Walker *et al*.^39^ used μCT BMDD data to evaluate the impact of age on bone in mice with osteocyte-specific deletion of Socs3 and/or Il6st. But rather than calculating the full distribution and using the parameters to define BMDD, they divided cortical BMDD into tertiles and so their approach didn’t use the full power of the method. Building upon these earlier studies, we now clearly demonstrate that μCT-based BMDD, an analysis that is available from scans generated by commonly used bone-phenotyping tool, can be utilized for assessing various aspects of MP. This expands the information available for assessing bone mineralization from tools like histomorphometry and avoids the specialized equipment and biopsies necessary to implement BSEM-based BMDD.

Despite the strengths of our study design and the quality of our data, there were several potential weaknesses in our approach. First, when Mashiatulla *et al.*^7^ compared μCT-based and BSEM-based BMDD data, they found that 6 μCT-based and BSEM-based BMDD parameters were highly correlated in HA phantoms (r^2^≥0.92), but in bone samples, two heterogeneity parameters (FWHM, coefficient of variation) from the two methods were not (r^2^≤0.05). They evaluated and eliminated two working hypotheses for this discrepancy: i.e. problems with beam hardening or partial volume effect caused by differences in porosity in the bone samples. They also suggested that a narrow range of heterogenicity values for μCT-based BMDD could account for this issue, but their data doesn’t support this view. A subsequent μCT-based BMDD paper using mice did not discuss this issue,^8^ while a μCT-based BMDD paper using human samples speculated, but did not test, that issues related to scan resolution or differences resulting from 2D and 3D images might be important.^9^ However, like we report here, these two papers reported treatment differences in heterogeneity parameters in μCT-based BMDD (measured by entropy or FWHM). Thus, we believe that the issue identified by Mashiatulla *et al.* is not a concern for μCT-based BMDD. Another issue is that in μCT-based BMDD, the left side of the curve does not reach to zero and has a long tail. Based on the binary thresholding approach used for traditional μCT phenotypes (where these values are viewed as “bone”), the low grayscale information represents under-mineralized bone. Others have adjusted the distribution to eliminate this issue.^6,37,40,41^ However, we think this adjustment discards relevant information about existing bone in μCT-based BMDD. Still, when a bone has an extreme phenotype, like the BMDD of VDR KO mice, users must be cautious, since this might impact FWHM and variance calculations. Finally, we used scans from different experiments that were generated with different μCT settings and reconstructed with different thresholds. Because of that, while we can still reliably compare samples within experiments, the methodological variability prevents direct comparisons across experiments. The use of HA phantoms to directly relate grayscale values to HA concentrations should be used to allow for comparisons of BMDD data across experiments and those generated with different μCT instruments.

This work lays the groundwork for research that furthers our understanding of mineralization. While we only assessed BMDD of cortical bone, bone mineralization and the MP are also important for trabecular bone. Human and mice studies have examined trabecular BMDD using BSEM^29,40,42^, and one human study has examined trabecular BMDD using μCT.^9^ We are currently exploring how to optimize μCT for assessing BMDD in mouse trabecular bone. The μCT-based BMDD method can also be used to generate information about primary and secondary mineralization. Ruffoni *et al.*^11^ mathematically modeled primary and secondary mineralization rates from a BSEM-based BMDD analysis of human trabecular bones. This has not yet been validated for μCT-based BMDD analysis, but if the equations are applicable, this analysis will allow us to test biological mechanisms controlling primary or secondary mineralization. Finally, *in vivo* μCT-based BMDD can be used to directly assess mineralization kinetics in longitudinal studies. This was previously demonstrated by Lukas *et al*.^40^ who examined the impact of mechanical loading on vertebral, trabecular bone mineralization kinetics in 15-week-old female mice.

In summary, here we provide important new data that shows μCT-based BMDD analysis is a valuable tool for the assessment of critical parameters that relate to MP. This technique provides novel information about the degree and heterogeneity of mineralization that are not available from other common bone phenotyping tools. Finally, the BMDD parameters can easily be calculated from existing μCT scans and so this data expands the utility of a common bone phenotypic tool while also increasing the ability to draw biological inferences about the nature of bone and the processes controlling bone mass through mineralization.

## Materials and Methods

### Animal Models

<u>(1) VDR KO mice:</u>^43^ Homozygous VDR KO mice^44^ and WT were fed an AIN93G-based rescue diet containing 20 g/kg calcium, 12.5 g/kg phosphorus, 200 g/kg lactose, and 1000 IU/kg Vitamin D_3_ (Research Diets, New Brunswick NJ, USA) and femora were harvest at 12 weeks of age and prepared for μCT analysis. <u>(2) Experimentally induced CKD and aging mice:</u>^13^ 16- and 78-wk-old C57BL/6J mice (Jackson Laboratory, Bar Harbor, ME, USA) were fed casein-based diets with 0.6% Ca and 0.9% P with/without 0.2% adenine (AD; Envigo-Teklad Diets, Madison, WI, USA) to induce CKD. After 6 wks on adenine/control diet, all mice were fed the control casein diet for 2 more weeks. Femora were harvested for μCT. <u>(3) Lrp5^A214V^ mice:</u>^16^ Male Lrp5^A214V^ and littermate control mice (WT) were fed purified AIN93G diets containing 0.4 % P and 1000 IU vitamin D/kg and 0.5% Ca (Research Diets, New Brunswick, NJ, USA) from weaning until 12 weeks of age. Femora were harvested for μCT analysis. <u>(4) iPTH and vehicle mice</u>: Male 19-wk-old C57BL/6NHsd mice (Envigo, Madison, WI, USA) were injected with vehicle or 100 ng/g/day of PTH (1-34) for 28 days. Tibiae were collected for μCT analysis. <u>(5) Genetically diverse mice fed normal and low calcium diets:</u>^19^ Male mice from 11 different inbred strains were fed either a 0.5% Ca (adequate) or 0.25% Ca (low) diet (AIN93G base with 200 IU vitamin D3/kg diet, Research Diets, New Brunswick, NJ, USA) starting at 4 wks of age. At 12 wks of age, femora were collected for μCT. All animal protocols were approved by the Institutional Animal Care and Use Committee at Indiana University School of Medicine or at Purdue University, and animal care was carried out in accordance with institutional guidelines.

### μCT Scans and Analysis

Because we used mice from a variety of experiments conducted at different times and in different research labs, the bone scanning methods were similar, but not identical. Details for each experiment are: <u>(1) VDR KO mice:</u> The femur midshaft was identified and a 240 µm distance was scanned (120 um above and below the midline) using a SkyScan 1276 with settings: 61 kV, 100 µA, 6 µm voxel size, 0.3 rotation step, frame averaging of 2, 0.5 aluminum filter with 360 rotation, 1500 ms exposure, and analyzed at 12 μm voxel size. <u>(2) CKD and aging study:</u> distal femora were scanned with 60 kV, 0.5 aluminum filter, frame average of 2, 0.7 rotation steps, and 8 µm voxel size.^13^ The scans were performed in SkyScan 1172, and 240 µm was analyzed in the region 3 mm proximal to distal growth plate. <u>(3) Lrp5^A214V^ mice:</u> femurs were scanned at the isotropic voxel size of 16 μm using an energy level of 55 kVp, an integration time of 300 ms, and an X-ray tube current of 145 μA in ScanCo μCT 40 system. 240 µm in midshaft was analyzed.^16^ <u>(4) iPTH treatment</u>: Tibial cortical bone in a 32 µm section 3 mm distal from the growth plate in the proximal tibia was scanned using a SkyScan 1172 with settings: 59 kV, 167 uA, 7.93 µm voxel size, 0.7 rotation. <u>(5) Genetically diverse mice fed adequate and low calcium diets:</u> femurs were scanned at the isotropic voxel size of 16 μm using an energy level of 55 kVp, an integration time of 300 ms, and an X-ray tube current of 145 μA in ScanCo μCT 40 system. 240 µm in midshaft was analyzed.^45^

### Data Processing for BMDD Analysis

All the samples for BMDD analysis were processed following randomization; investigators assessing the experimental outcomes were blinded to the treatment or stain information. ISQ files of bone scans from ScanCo were converted from 16-bit to 8-bit format and BMP files from SkyScan were opened in the CTAn 1.21.1 micro-CT software (Bruker). During conversion or reconstruction, thresholds for defining bone were reselected to ensure that the histograms were not saturated. If histograms were saturated previously scanned bones were reconstructed again. In the iPTH study, there was a subset of bone scans that were oversaturated but the bones from this study were not available for re-scanning so the highest density pixels were lost in these bones.

Within each experiment, bones from each group were opened to identify lowest grayvalue in the image that could be classified as bone. To do this, the user needed to switch previews between the raw image and the binary-selection images as they determined the grayvalue that was still considered bone. Once determined, the low threshold value was used for all samples within an experiment (Supplemental Fig. 1). CTAn provides a value for the total number of voxels above the lower threshold and then uses data from each voxel to calculate the bone volume at each greyscale value as well as the percentile of bone volume present at each greyscale value (i.e. grayscale volume normalized to total volume). These values were plotted to define the BMDD for each sample. The number of voxels at each greyscale (i.e. the raw data) was calculated by multiplying percentage of bone volume at each grayscale by the total number of voxels in the scan. Eleven parameters from the BMDD were calculated for each individual sample using RStudio (Appendix 1. BMDD Calculations): total volume (or voxels) in the distribution, lower 5% of the distribution, upper 5%, the maximum greyscale value (Max), center (aka the mode), peak height, full width at half maximal peak height (FWHM), mean degree of mineralized bone (MDMB), variance, skewness, and kurtosis.

### Statistics

These experiments were designed to have the statistical power to see treatment/strain differences using traditional μCT or DEXA phenotypes. All experimental results represent the data collected from individual mice. Outliers were identified as values whose z-score was in the extreme 2.5% tails of the line/diet group and values outside this range were removed were removed prior to analysis.

*A) Evaluating Differences between groups:* For comparing BMDD data from the preclinical models with 2 groups, we used 2-tailed Student’s t-test and conducted this analysis in Excel. All the means, standard errors (SEM) and the number of samples used for each analysis are reported in Supplemental Table 1.

The study on the 11 inbred mouse strains was a factorial design so for the examination of the BMDD data from this study. We conducted 2-way ANOVA analysis using SAS Enterprise Guide 8.3 (SAS Institute Inc., Cary, NC, USA). Prior to analysis, the covariate effect of body weight (BW) and femur length (FL) on all parameters were determined by Pearson’s correlation and when significant, added as a covariate to the analysis. A normal distribution of the residuals was determined by assessing Q-Q plots, Predicted vs Residual plots, and with the Shapiro-Wilk test. Equality of variance was tested by assessing Predicted vs Residual plots and by Brown-Forsythe test for ANOVA in the residual data, and ANCOVA in body size corrected residual data. Transformations performed are in Supplemental Table 2. Two-way ANOVA or ANCOVA was used to test for the significant main effects (G, D) and a GxD interaction. The p-values for main effects and the interaction are reported in Supplemental Table 2. When significant interactions were observed, the Tukey-Kramer post-hoc test was used to determine differences among the groups. All the means, standard errors, and the number of samples used for each analysis for statistics for this study are reported in Supplemental Table 3.

*B) Correlations and PCA:* Pearson’s correlations and PCA were performed in Origin 2024 (OriginLab, Northampton, MA, USA). The number of PC considered significant was determined by identifying the ‘elbow’ of the Scree plot.

*C) Multiple Linear Regressions:* Analysis was performed on data from the 11 genetically diverse mouse lines in Origin 2024 to assess the combination of bone parameters that best predicted US 1^st^. Backward selection model was performed with all the BMDD parameters as well as BMD, BMC, and Ct.Th. The r^2^ values were adjusted for the number of independent variables in the model.

Methods for Biodent and 4-point bending performed are in Supplemental Materials.

## Data Availability Statement

The data that support the findings of this study are available upon request from the corresponding author.

## Acknowledgments

This work was supported by start-up funds from the University of Texas at Austin to JCF and by the National Institutes of Health NIH-NIAMS (5T32AR065971-06 to SUO). We thank Drs. Corinne Metzger and Matthew Allen for providing data from the ageing and CKD study.

## Contributions

Serra Ucer Ozgurel: Conceptualization, Methodology, Software, Validation, Formal Analysis, Investigation, Resources, Visualization, Data Curation, Writing – Original Draft, Writing – Review & Editing; Arish Maredia: Methodology, Software, Writing – Review & Editing ; Joseph Sheeran: Formal analysis, Writing – Review & Editing; Laurent Marichal: Methodology, Formal Analysis, Writing – Review & Editing; James Fleet: Conceptualization, Methodology, Project administration, Investigation, Resources, Writing – Original Draft, Writing – Review & Editing, Supervision, Funding acquisition.

## Appendix 1. BMDD Calculations

| BMDD Parameters | Mathematical Definition/Equation | Mathematical Meaning | Biological Meaning |
| --- | --- | --- | --- |
| Height | Max (number of voxels*normalized data) = max (raw data) | Highest y value of the distribution | Height reflects the amount of bone at the most frequent mineral level |
| Number of Voxels | $(1 \text{ mm}/\text{resolution})^3$ | Summation of pixel in the selected region | Indicator of amount of bone, independent of amount of mineral |
| Maximum Grayvalue | Highest x with nonzero data | Highest x value with a raw data | Highest mineralization level presented in a given bone |
| Mean Degree of Mineralized Bone (MDMB) | $\sum_{x_i}^{255} (\text{raw data})/n$ | Each voxel is calculated and MDMB is calculated with the consideration of the mineral levels | Defines the average mineral level |
| Center | X value in the highest y value | x corresponding to highest value of y | Most common mineral level present in a given bone |
| Full Width Half Maximum | x length in ½ max raw data | Mineral level width, which covers the half volume of bone | Indicator for heterogeneity |
| Upper 5% (u) | x value for 95% of raw data | x value where 95% of the bone is covered | Highest 5% mineralization level present in a given bone |
| Lower 5% | x value for 5% of raw data | x value where 5% of the bone is covered | Lowest 5% mineralization level present in a given bone |
| Variance ($s^2$) | $(\sum f_i (X_i - \text{MDMB})^2)/n-1$ | The average of squared standard deviations from the mean | The heterogeneity of distribution, the spread of data |
| Skewness | $\frac{\sqrt{n(n-1)}}{n-2} * \frac{(\sum f_i (x_i - \text{MDMB})^3)/n}{(\text{Variance})^{3/2}}$ | Measurement of asymmetry in a distribution | The asymmetry of the distribution. Eg. Positive skew: majority of a given bone is less mineralized |
| Kurtosis (kurt) | $\frac{\sum f_i (x_i - \text{MDMB})^4}{s^2 * n}$ | Tailedness of the distribution | Leptokurtic BMDD indicates narrower peak and fatter tails, indicating overall more heterogenicity |

