## Supplemental Methods and Figures for "μCT-based, Three-Dimensional Cortical Bone Mineral Density Distribution reveals Unique Phenotypes related to Bone Material Properties"

**Supplemental Materials and Methods:**

*Biodent:* Reference point indentation (RPI) was conducted using Biodent$^{TM}$ (Active Life Scientific, Inc., Santa Barbara, CA, USA) on femora from 11 genetically diverse inbred mouse strains; there were the same bones that were scanned for μCT. A mouse-specific probe (PB3) was used and the diaphysis of the femur was placed parallel to the center rod with the distal femur side being on the left. The test was applied to the midshaft of the bone with no preconditioning, 10 indentation cycles performed at 2 Hz, with a force of 2 N. A minimum 6 measurements were obtained per bone. Three measurements were conducted with a poly (methyl methacrylate) reference material before and after each assessment session and these values were used to normalize measurements across sessions. Nine parameters were generated from the RPI analysis: 1st cycle indentation distance (ID 1st), 1st cycle unloading slope (US 1st), 1st cycle creep indentation distance (CID 1st), 1st Total Indentation Distance (T1D 1st). Indentation Distance Increase (IDI 1stL), Average Creep Indentation Distance (Ave CID1stL), Average energy dissipated (Ave ED 3rd-L), Average unloading slope (Ave US1stL), and average loading slope (Ave LS1stL).

*4-point bending:* Four-point bending was performed in tibiae collected from the CKD and Aging study.[14] Force-displacement curves were produced in bones that were loaded to failure using an ElectroForce mechanical testing instrument and a 0.025 mm/s rate (TA Instruments, New Castle DE, USA). Structural mechanical properties were obtained by utilizing a custom MATLAB code and MP were obtained from standard beam-bending equations, i.e. ultimate force, total displacement, stiffness, total work, ultimate stress, modulus, resilience and toughness were obtained.

**Supplemental Figures:**

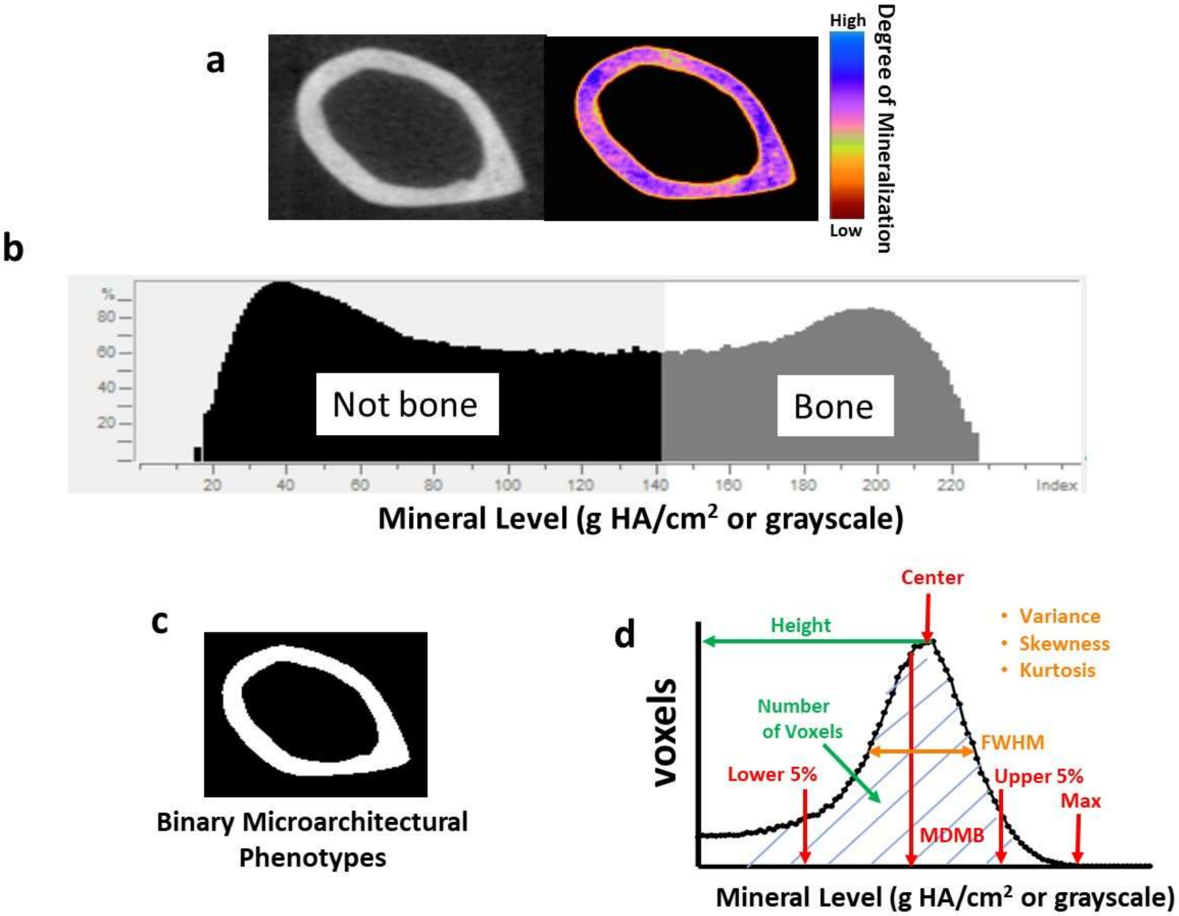


**Supplemental Figure 1. An Overview of BMDD Analysis from µCT. (A)** µCT scan of cortical bone in grayscale and as a false color image that reflects different mineral densities based on the color scale to the right. **(B)** The distribution of densities from a cortical bone scan. **(C)** A "bone/not bone" image resulting from thresholding is used to calculate traditional µCT cortical endpoints. **(D)** A BMDD from a distribution of densities and the parameters that describe the distribution. Images are obtained from CTAn.

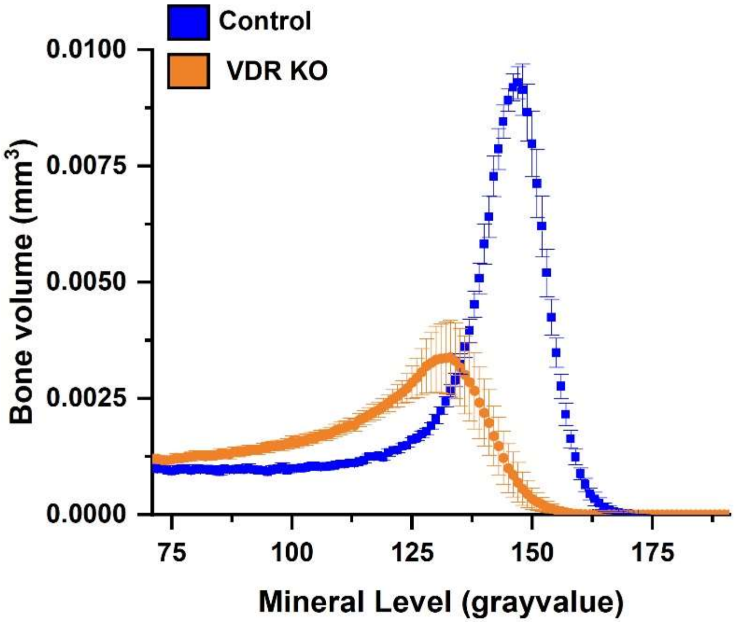


**Supplemental Figure 2. The Impact of VDR KO on the Raw Bone Volume BMDD in the Femoral Cortical Bone Midshaft:** The raw bone volume BMDD curve of the femoral cortical midshaft from 12-wk old vitamin D receptor knockout (VDR KO n=12) and littermate control (n=5) mice fed 0.5% Ca AIN93G diets. BMDD are plotted as mean±SEM. The normalized % bone volume data from these mice are presented in Figure 2.

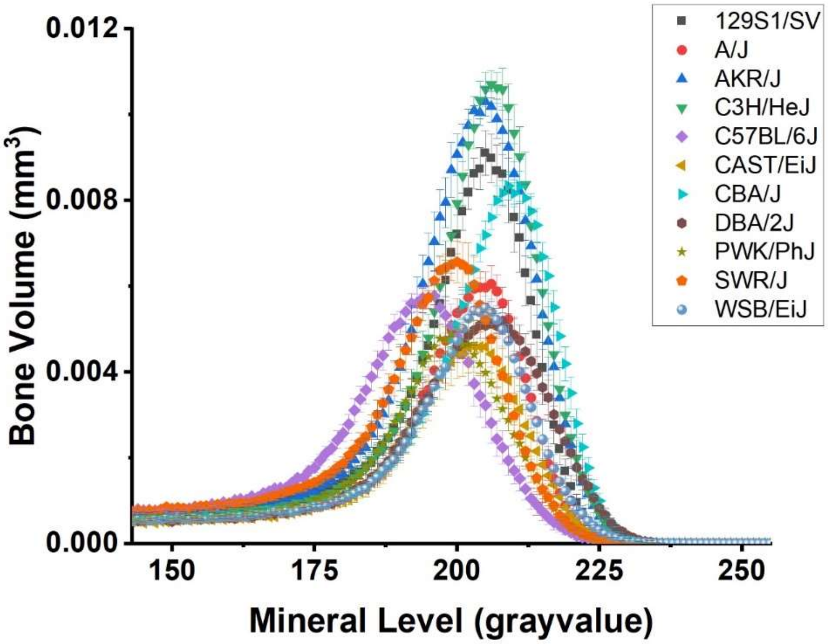


**Supplemental Fig 3. The Impact of Natural Genetic Variation on the Raw Bone Volume BMDD in the Femoral Cortical Bone Midshaft:** The raw bone volume BMDD curve of the femoral cortical midshaft from 12-wk old mice fed 0.5% Ca AIN93G diets. Data are from 11 genetically distinct inbred mouse lines (n=6-8 mice per line). BMDD are plotted as mean±SEM. The normalized % bone volume data from these mice are presented in Figure 3.

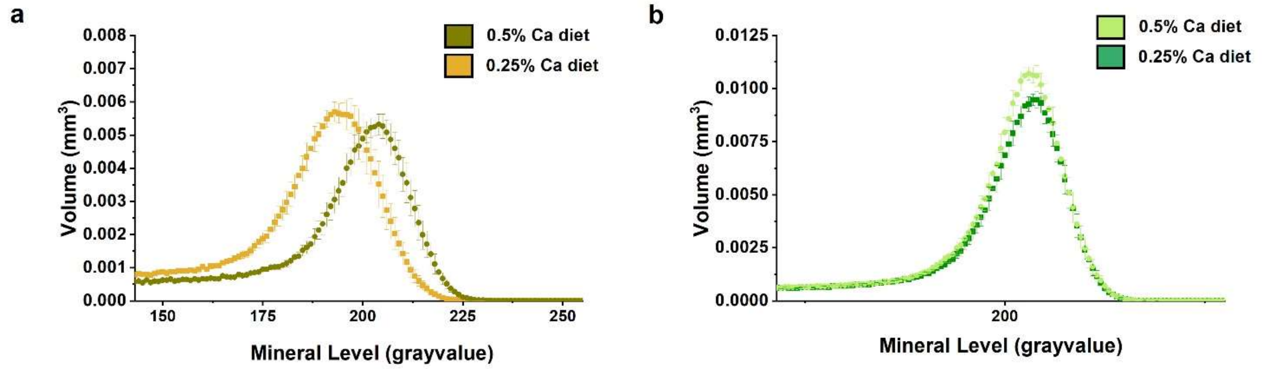


**Supplemental Figure 4. The Impact of Gene x Diet on the Raw Bone Volume BMDD in the Femoral Cortical Bone Midshaft:** The raw bone volume BMDD curve of the femoral cortical midshaft from 12-wk old mice fed 0.5% Ca or 0.25% Ca AIN93G diets. Data are from **A.** CAST/EiJ and **B.** C3H/HeJ mice (n=7-8 mice per diet per line). BMDD are plotted as mean±SEM. The normalized % bone volume data from these mice are presented in Figure 4C and 4D.

a b

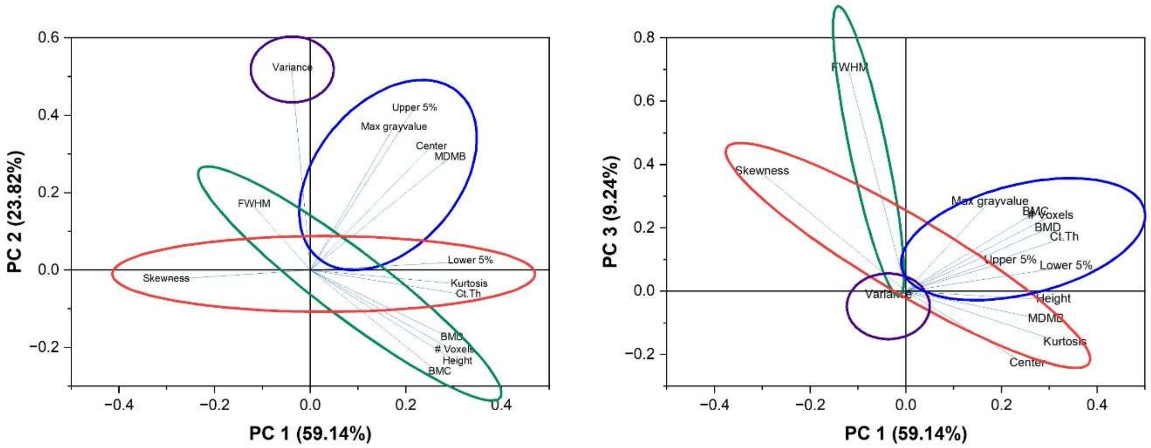


c

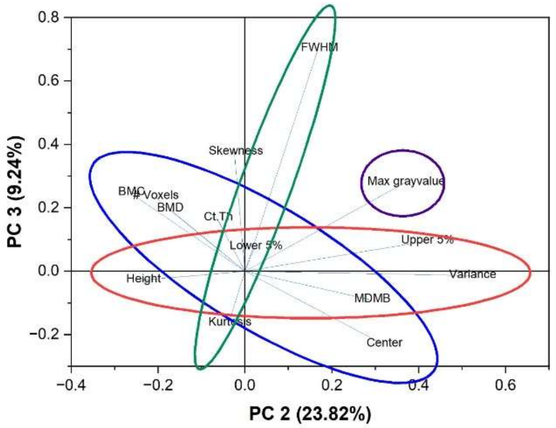


d

| | PC1 | PC2 | PC3 | PC4 |
|---|---|---|---|---|
| **Variance explained (%)** | 59.14 | 23.82 | 9.24 | 3.45 |
| **BMD** | 0.2937 | -0.1755 | 0.1969 | 0.2946 |
| **BMC** | 0.2681 | -0.2644 | 0.2473 | 0.2507 |
| **Ct Th** | 0.3281 | -0.06467 | 0.1677 | -0.1586 |
| **Total Voxels** | 0.2997 | -0.2084 | 0.2360 | 0.1157 |
| **Height** | 0.3053 | -0.2368 | -0.0281 | 0.1603 |
| **Maximum gv** | 0.1730 | 0.3676 | 0.2803 | 0.4321 |
| **Lower 5% Cutoff** | 0.3324 | 0.02197 | 0.07710 | -0.3104 |
| **Upper 5% Cutoff** | 0.2157 | 0.4168 | 0.09491 | 0.00674 |
| **Center** | 0.2506 | 0.3174 | -0.2297 | -0.06281 |
| **MDMB** | 0.2899 | 0.2898 | -0.0900 | -0.1213 |
| **Kurtosis** | 0.3300 | -0.03774 | -0.1634 | -0.2664 |
| **Skewness** | -0.3040 | -0.02456 | 0.3757 | 0.2327 |
| **Variance** | -0.03961 | 0.5207 | -0.0136 | 0.2949 |
| **FWHM** | -0.1205 | 0.1683 | 0.7021 | -0.5218 |

**Supplemental Figure 5: 2D PCA Plots from 3D Plotted in Figure 6**. **A.** PC1 and PC2 **B.** PC1 and PC3 **C.** PC2 and PC3. **D.** Table of Extracted eigenvectors for 4 PCs. Orange highlighted values are those that have a weigh in the PC greater than 0.289. Colors in PC graphs are used as a visual aid to identify subgroups of related phenotypes.
